# Dendritic *atoh1a+* cells serve as transient intermediates during zebrafish Merkel cell development and regeneration

**DOI:** 10.1101/2023.09.14.557830

**Authors:** Evan W. Craig, Erik C. Black, Camille E.A. Goo, Avery Angell Swearer, Nathaniel G. Yee, Jeffrey P. Rasmussen

## Abstract

Sensory cells often adopt specific morphologies that aid in the detection of external stimuli. Merkel cells encode gentle touch stimuli in vertebrate skin and adopt a reproducible shape characterized by spiky, actin-rich microvilli that emanate from the cell surface. The mechanism by which Merkel cells acquire this stereotyped morphology from basal keratinocyte progenitors is unknown. Here, we establish that dendritic Merkel cells (dMCs) express *atonal homolog 1a (atoh1a)*, extend dynamic filopodial processes, and arise in transient waves during zebrafish skin development and regeneration. We find that dMCs share molecular similarities with both basal keratinocytes and Merkel cells, yet display mesenchymal-like behaviors, including local cell motility and proliferation within the epidermis. Furthermore, dMCs can directly adopt the mature, microvilliated Merkel cell morphology through substantial remodeling of the actin cytoskeleton. Loss of Ectodysplasin A signaling alters the morphology of dMCs and Merkel cells within specific skin regions. Our results show that dMCs represent an intermediate state in the Merkel cell maturation program and identify Ectodysplasin A signaling as a key regulator of Merkel cell morphology.

## INTRODUCTION

Organ development and function require that constituent cells adopt precise shapes. For example, epithelial cells often develop elaborate actin-based membrane protrusions integral to organ function, such as the brush border of intestinal enterocytes or the stereocilia of inner ear hair cells (reviewed by Sharkova et al., 2023). Defects in the morphogenesis of actin-based protrusions are linked to a variety of diseases, including colorectal cancer and deafness. Thus, elucidating the mechanisms of how epithelial cells adopt specific shapes is relevant to understanding both the basis for organ function and the etiology of human pathologies.

Merkel cells (MCs) are mechanosensory epidermal cells that interact with somatosensory neurites to form the MC-neurite complex, which mediates detection of gentle touch stimuli (reviewed by Woo et al., 2015). MCs populate diverse vertebrate skin compartments, including human plantar and palmar skin, mouse hairy and glabrous skin, and teleost facial and trunk skin (reviewed by Hartschuh et al., 1986; Whitear, 1989). MCs have a remarkably consistent morphology across these diverse vertebrate skin types. The core morphological features of MCs include a small and spherical cell body, a high nuclear-to-cytoplasmic ratio, and the presence of neurosecretory granules (Hartschuh et al., 1986; Whitear, 1989). But perhaps the most striking morphological feature of MCs are the numerous actin-rich microvilli that emanate off the cell surface, giving MCs a “mace-like” morphology (Hartschuh et al., 1986; Lane and Whitear, 1977; Sekerková et al., 2004; Takahashi-Iwanaga, 2003; Takahashi-Iwanaga and Abe, 2001; Toyoshima et al., 1998; Whitear and Lane, 1981).

The precise and reproducible cell shape of MCs has been proposed to impact their sensory function (Garant et al., 1980; Takahashi-Iwanaga, 2003; Takahashi-Iwanaga and Abe, 2001; Toyoshima et al., 1998; Yamashita et al., 1993); however, it is unknown how MCs acquire this stereotyped morphology. Cre-based lineage tracing experiments indicate that MCs derive from basal keratinocyte precursors in both mammalian and zebrafish skin (Brown et al., 2023; Morrison et al., 2009; Van Keymeulen et al., 2009). While these experiments begin to address MC progenitor identity, they fail to shed light on how cells within this lineage lose basal keratinocyte characteristics and adopt the hallmark MC morphology. A deeper understanding of the MC lineage may shed light on the origins of Merkel cell carcinoma, an aggressive skin cancer of unclear cellular origin (reviewed by Becker et al., 2017; Harms et al., 2018).

Interestingly, several histological observations support the presence of cellular intermediates in the MC lineage. First, ultrastructural studies identified “transitional cells” that exhibit characteristics of both keratinocytes and MCs, including fine cytoplasmic filaments and dense-core granules, respectively (English, 1974; English, 1977; Garant et al., 1980; Tachibana and Ishizeki, 1981; Tachibana and Nawa, 1980). Second, immunostaining for two MC markers— cytokeratins 18 and 20—revealed cells with oblong or dendritic morphologies in developing human plantar skin, human and rodent oral mucosa, and murine touch domes (Kim and Holbrook, 1995; Moll et al., 1984; Nakafusa et al., 2006; Tachibana et al., 1997; Tachibana et al., 1998). Although these heterogeneously shaped cells have been referred to as dendritic MCs (dMCs), it is unknown whether dMCs represent a developmentally immature MC state or a functionally distinct subpopulation of neuroendocrine cells (reviewed by Xiao et al., 2014). Live-cell imaging of MC development would be the most direct way to distinguish these possibilities. However, to date, technical limitations of mammalian systems have precluded visualization of dMC or MC dynamics in living animals.

We recently established zebrafish as an *in vivo* model for MC studies (Brown et al., 2023). Here, we leverage the optical accessibility of zebrafish skin to directly visualize individual cell behaviors during the MC maturation process. By visualizing key stages of skin organogenesis and regeneration using an F-actin reporter expressed in MCs, we describe a morphologically distinct population of epidermal cells that shares characteristics with transitional cells and dMCs observed in other types of vertebrate skin. Importantly, we document the direct maturation of dMCs into MCs through cytoskeletal rearrangements. Furthermore, we show that Ectodysplasin A (Eda) signaling is required for MCs to develop microvilli within trunk skin. Together, our results provide the first *in vivo* characterizations of MC precursor states and identify Eda signaling as a key regulator of zebrafish MC morphogenesis.

## RESULTS

### A transient, morphologically distinct population of keratinocyte-derived *atoh1a+* cells emerges during skin development

Staging of zebrafish post-embryonic development relies on standard length (SL) in millimeters, with sexual maturity attained ∼18 mm SL (Parichy et al., 2009). Cells with the prototypical MC morphology—spherical with actin-rich, microvillar protrusions of ∼1-2 microns in length— populate several adult zebrafish skin compartments, including trunk skin (**Figure 1A,B**; (Brown et al., 2023)). *Atonal homolog 1* (*Atoh1*) encodes a transcription factor necessary and sufficient for murine MC development (Morrison et al., 2009; Ostrowski et al., 2015; Van Keymeulen et al., 2009). Zebrafish *atoh1a* is an ortholog of murine *Atoh1*, and zebrafish MCs express reporters inserted into the upstream region of the *atoh1a* locus (Brown et al., 2023). Using a nuclear localized *atoh1a* reporter to label MCs, we previously found that trunk MCs begin to appear during the onset of squamation (scale formation) at approximately 9 mm SL and rapidly increase in density from 10-15 mm SL (Brown et al., 2023). As this nuclear reporter did not allow for visualization of cell shapes, it is unclear whether morphological heterogeneity exists within developing zebrafish MCs, including the possible presence of dMCs.

**Figure 1.**
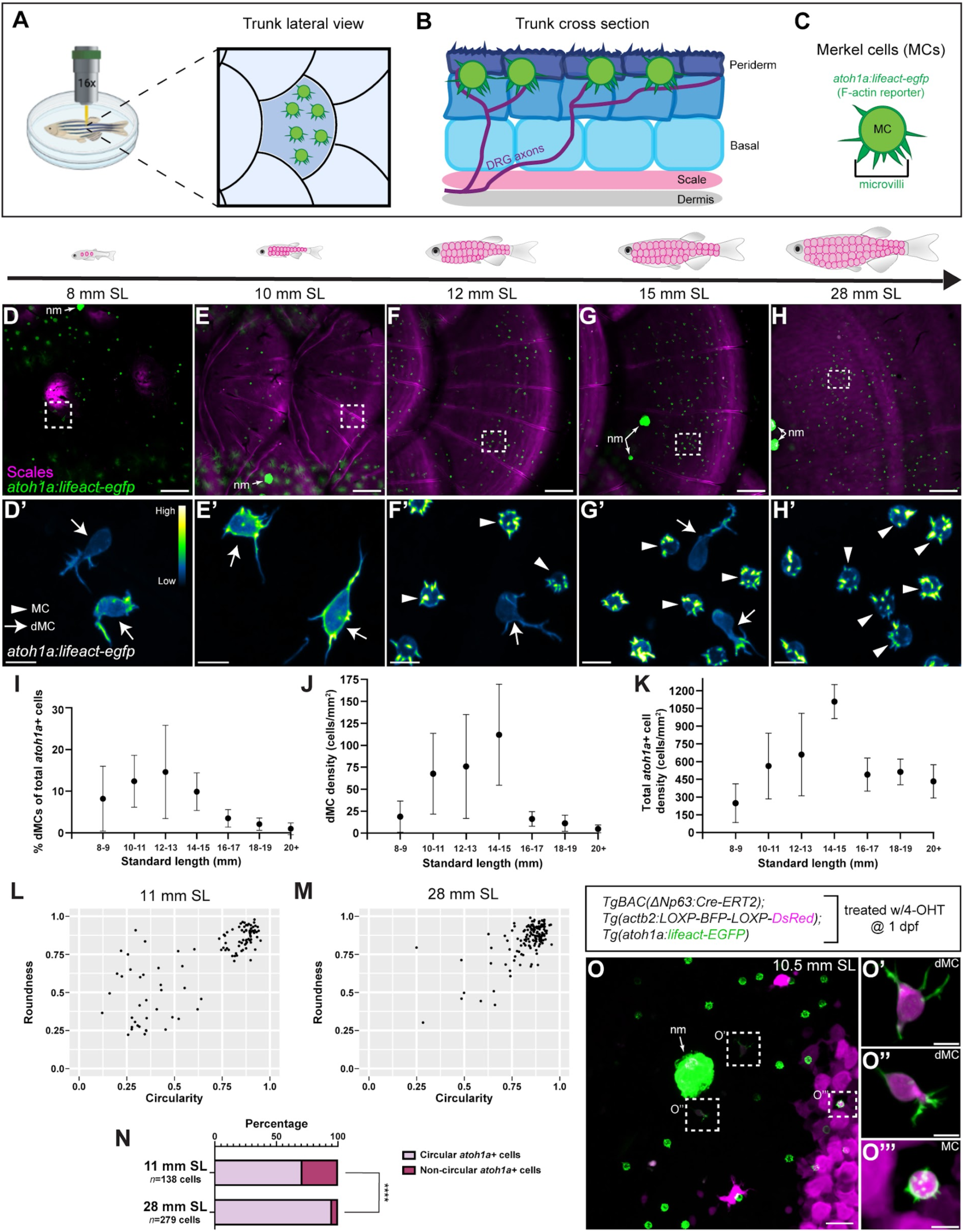
Identification of a transient, morphologically distinct population of keratinocyte-derived *atoh1a*+ cells during skin development. **(A)** Schematic depicting live-imaging methodology for visualizing MCs (green) within the epidermis atop scales along the lateral zebrafish trunk skin. **(B)** Illustration showing organization of adult zebrafish trunk skin in cross section. Note the stratified layers of keratinocytes above the bony scale. Merkel cells are shown in green near the outermost epidermal layer. **(C)** The *Tg(atoh1a:lifeact-egfp)* reporter allows visualization of the MC F-actin cytoskeleton, including short and spiky microvilli structures. **(D-H)** Representative confocal micrographs of *atoh1a+* MCs (green; *Tg(atoh1a:lifeact-egfp)*) and scales (magenta; Alizarin Red S staining) along the lateral trunk in juvenile to adult zebrafish of the indicated stages. White dashed boxes indicate regions magnified in (D’-H’). **(D’-H’)** *Tg(atoh1a:lifeact-egfp)* signal intensity is color-coded using the “Green Fire Blue” lookup table. MCs denoted by arrowheads, dMCs denoted by arrows. **(I)** The percentage of *atoh1a+* cells with dMC morphology out of total number of *atoh1a+* cells plotted against SL. Each dot represents the mean dMC percentage calculated from at least *n*=3 animals of that size range. For each individual zebrafish, multiple confocal micrographs were acquired from the trunk region at 1.5x magnification and *atoh1a+* cells were manually categorized as “MC” or “dMC” based on morphology and grouped for input data to calculate the dMC ratio shown. **(J)** Plot of dMC density relative to SL from the dataset analyzed in (I). **(K)** Plot of total *atoh1a+* cell density (dMCs and MCs) relative to SL from the dataset analyzed in (I). **(L,M)** *atoh1a+* cell shape analysis for animals of the indicated stages. Black dots represent individual *atoh1a+* cells that were assigned values for circularity and roundness after confocal micrographs of *Tg(atoh1a:lifeact-egfp)* were thresholded to acquire *atoh1a+* cell shape outlines. Top right quadrant indicates cells with the most round and circular morphologies, indicative of mature MCs. Cells falling outside of the top right quadrant possess less round and circular morphologies, indicative of dMCs. *n*=100 randomly selected cells displayed for each stage. **(N)** Stacked barcharts of *atoh1a+* cell shape analysis performed in L-M. *atoh1a+* cells in adult skin have significantly more circular morphologies than *atoh1a+* cells in juvenile skin (****, P<0.0001; Fisher’s Exact Test). **(O-O’’’)** Confocal micrographs of the trunk epidermis from a *TgBAC(ΔNp63:Cre-ERT2); Tg(actb2:LOXP-BFP-LOXP-DsRed); Tg(atoh1a:lifeact-egfp)* juvenile treated with 4-OHT at 1 dpf. Note the mosaic DsRed expression (magenta) in basal keratinocytes and derivatives. DsRed+ dMCs shown in O’-O’’, along with a DsRed+ MC in O’’’ indicate both MC-types derive from a shared basal keratinocyte progenitor population. nm, neuromasts containing clusters of *atoh1a+* hair cells. Error bars in (I,J,K) display SD of mean. Scale bars: 50 µm (D-H), 20 µm (O), and 5 µm (D’-H’,O’-O’’’).

To assess MC morphology during squamation, we conducted a similar developmental staging time course using *Tg(atoh1a:lifeact-egfp)*, which expresses a filamentous actin (F-actin) reporter in MCs, allowing detailed visualization of the MC actin cytoskeleton *in vivo* (**Figure 1C**; (Brown et al., 2023)). To simultaneously visualize scales, we stained animals with Alizarin Red S, which labels the calcified scale matrix. Like our previous observations in adults, we identified cells with the prototypical MC morphology at late juvenile and adult stages (**Figure 1F-H**, arrowheads). Interestingly, we also observed a second population of *atoh1a+* cells with weaker transgene signal and highly variable morphologies, characterized by ovoid cell bodies and long, filopodial-like actin-rich protrusions (**Figure 1D-G**, arrows). For consistency with the literature (Boot et al., 1992; Kim and Holbrook, 1995; Moll et al., 1984; Nakafusa et al., 2006; Tachibana et al., 1997; Tachibana et al., 1998), we hereafter refer to these cells as dMCs, although we note that whether dMCs are related to MCs or represent an alternative fate has not been established. Manual cell counting from our imaging dataset revealed dMCs appeared in highest frequency and density at 10-15 mm SL, when scales expand and total MC density rapidly increases (**Figure 1I-K**). Next, we quantified *atoh1a* cell shapes at two different developmental stages by assigning circularity and roundness values to thresholded images from our dataset. We defined “circular” cells as those having circularity and roundness values >0.7. We found that 95.3% of *atoh1a+* cells in adult skin fell into the circular category, likely representing the mature, microvilliated MC type (**Figure 1M,N**). By contrast, only 71.1% of *atoh1a*+ cells in juvenile skin fell into the circular category, with numerous *atoh1a*+ cells adopting more oblong and less round shapes. (**Figure 1L,N**). Thus, our results indicate that a morphologically distinct population of *atoh1a+* cells transiently populates the trunk skin during squamation.

Using a tamoxifen-inducible Cre driver expressed in basal keratinocytes [*TgBAC(ΔNp63:Cre-ERT2)*; (Brown et al., 2023)] and a quasi-ubiquituous Cre reporter transgene [*Tg(actb2:LOXP-BFP-LOXP-DsRed)*; (Kobayashi et al., 2014)], we previously found that zebrafish MCs derive from *ΔNp63*-expressing basal keratinocytes (Brown et al., 2023). To determine if basal keratinocytes also give rise to dMCs, we treated *TgBAC(ΔNp63:Cre-ERT2); Tg(actb2:LOXP-BFP-LOXP-DsRed); Tg(atoh1a:lifeact-egfp)* embryos with 4-OHT to induce Cre-ERT2 activity at 1 day post-fertilization (dpf). This resulted in permanent DsRed expression in basal keratinocytes and their cellular derivatives. We then raised animals to squamation stages. Due to transgene mosaicism and/or incomplete Cre-ERT2 activation, not all basal keratinocytes expressed DsRed (**Figure 1O**). Along with DsRed+ MCs (**Figure 1O’’’**), we observed that a subset of dMCs in juvenile skin were DsRed+ (**Figure 1O’-O’’**). Interestingly, we often identified DsRed+ dMCs at a distance from the nearest DsRed+ basal keratinocyte (**Figure 1O**). These findings demonstrate a basal keratinocyte origin for dMCs and suggest they may be motile.

### dMCs are the primary *atoh1a*+ cell type during early stages of skin regeneration

Our identification of dMCs during squamation prompted us to hypothesize that dMCs may also appear during specific stages of scale regeneration. To induce scale regeneration, we performed scale “plucking” with fine forceps to remove both the bony scale and overlying epidermis containing MCs (**Figure 2A**). Scale removal triggers a wound healing response from neighboring keratinocytes, which migrate from surrounding, non-injured scales to cover the denuded area within hours (Richardson et al., 2016). Over the course of several days, keratinocytes then proliferate and reestablish a stratified epidermis (Richardson et al., 2016), while dermal osteoblasts proliferate and undergo hypertrophy to regenerate the bony scale (Aman et al., 2018; Cox et al., 2018; De Simone et al., 2021; Iwasaki et al., 2018; Sire et al., 1997).

**Figure 2.**
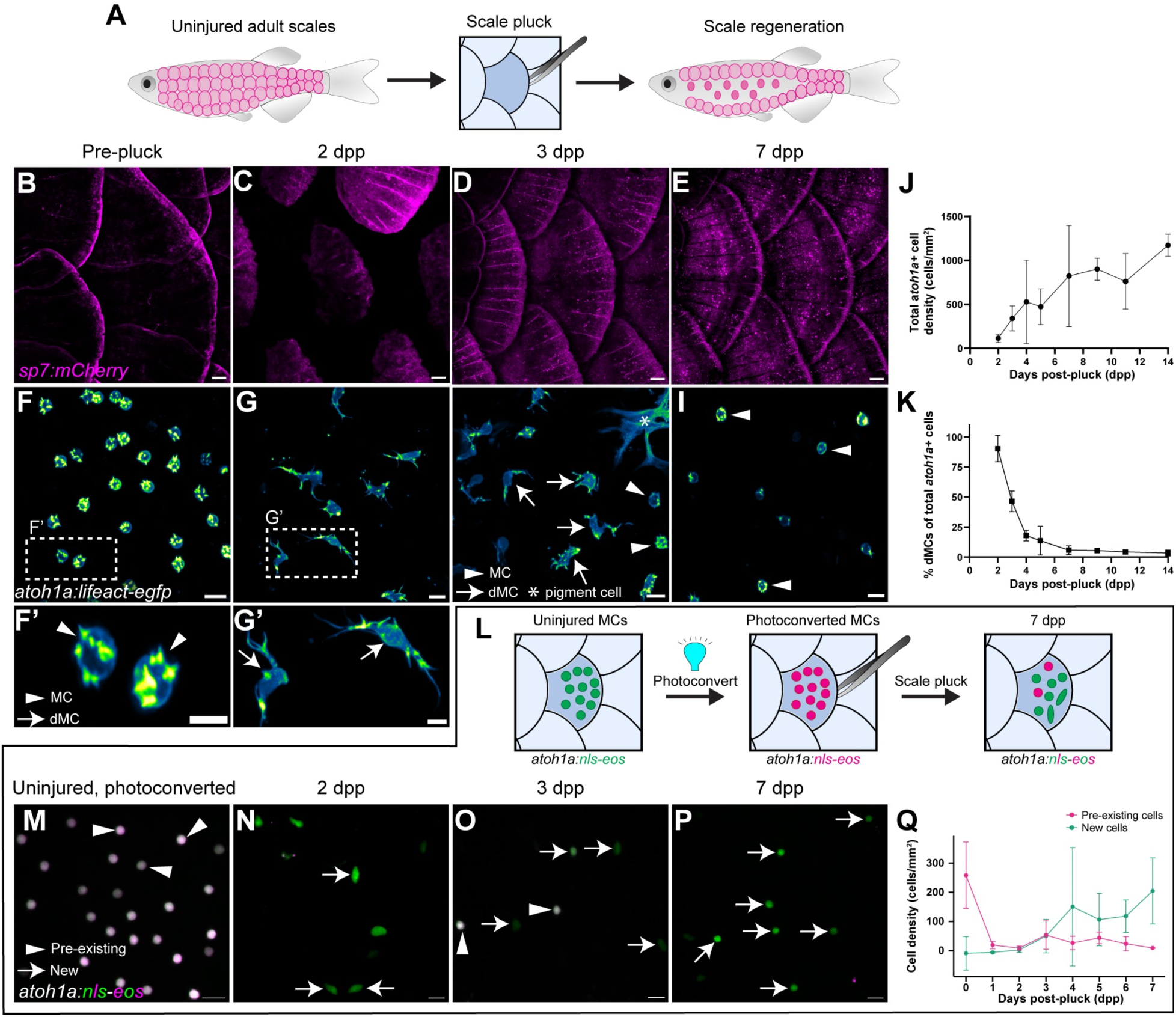
dMCs are the primary *atoh1a*+ cell type during the early stages of skin regeneration. **(A)** Illustration of the zebrafish scale pluck regeneration model. Forceps are used to physically “pluck” scales from the adult zebrafish trunk to induce injury and trigger regeneration of dermal scales and overlying epidermis. **(B-E)** Representative confocal micrographs of scale-forming osteoblasts (magenta; *Tg(sp7:mCherry)*) at the indicated stages of regeneration. **(F-I)** Representative confocal micrographs of *atoh1a+* cell shapes within the scale epidermis at the indicated stages of scale regeneration. Yellow arrows denote MCs, green arrows denote dMCs, white arrow denotes an autofluorescent pigment cell. White dashed boxes indicate regions magnified in (F’,G’). **(J)** Plot of total *atoh1a+* cell density (dMCs and MCs) over the course of scale regeneration. Each dot represents confocal images collected and analyzed from multiple zebrafish of the corresponding timepoint (2-7 dpp, *n*=8-13; 9-14 dpp, *n*=2-4). **(K)** Quantification of dMC frequency during scale regeneration. Each dot represents the mean dMC frequency from the dataset analyzed in (J). **(L)** Illustration depicting the design of the *atoh1a+* cell photoconversion experiment. *Tg(atoh1a:nls-Eos)* expresses a nuclear-localized and photoconvertible Eos protein in *atoh1a+* cells (dMCs and MCs). *atoh1a+* cells (green) in uninjured scales are irreversibly photoconverted from green to magenta using UV light. Scales containing photoconverted cells are then plucked to induce regeneration. Pre-existing *atoh1a+* cells will contain photoconverted nls-Eos (magenta) in the new scale region whereas new *atoh1a+* cells will contain only non-photoconverted nls-Eos (green). **(M-P)** Representative confocal micrographs of the photoconverted *Tg(atoh1a:nls-Eos)* scale epidermis pre-scale pluck (M) and post-pluck (N-P). Arrowheads denote pre-existing cells (containing photoconverted nls-Eos), arrows denote *de novo* generated cells (containing only non-photoconverted nls-Eos). **(Q)** Quantification of pre-existing (magenta line) and new (green line) *atoh1a+* cells from photoconversion experiment at the indicated stages of regeneration. Each dot represents the mean *atoh1a+* cell density from *n*=3-4 fish. Scale bars, 100 µm (B-E), 10 µm (F-I), 5 µm (F’, G’), and 10 µm (M-P).

To determine whether MCs populated the regenerating scale epidermis, we plucked scales from animals expressing *Tg(atoh1a:Lifeact-egfp)* and *Tg(sp7:mCherry)* (Singh et al., 2012), a reporter for scale-forming osteoblasts (**Figure 2A,B**). As expected, scales underwent substantial regeneration within the first 7 days post-pluck (dpp) (**Figure 2B-E**). *atoh1a+* cells appeared above regenerating scales beginning at 2 dpp and rapidly increased in density until 7 dpp, after which the density plateaued (**Figure 2F-J**). Strikingly, dMCs comprised ∼95% of the *atoh1a+* cells present at 2 dpp (**Figure 2G,G’,K**) and ∼45% of the *atoh1a+* cells at 3 dpp (**Figure 2H,K**). After 3 dpp, the proportion of dMCs gradually decreased, with MCs becoming the predominant *atoh1a*+ cell type at later stages of regeneration (**Figure 2I,K**)

dMCs and MCs on regenerating scales could arise from either *de novo* production or movement of pre-existing *atoh1a+* cells from surrounding, uninjured regions of epidermis. To distinguish between these possibilities, we irreversibly photoconverted *atoh1a+* cells expressing a nuclear localized version of the photoconvertible protein Eos [*Tg(atoh1a:nls-Eos);* (Pickett et al., 2018)] and then plucked scales (**Figure 2L,M**). In this experimental paradigm, pre-existing *atoh1a+* cells contain both photoconverted and non-photoconverted nls-Eos, whereas newly differentiating *atoh1a+* cells contain only non-photoconverted nls-Eos. Notably, at 2 and 3 dpp the epidermis contained many newly differentiated cells with oval nuclear morphologies that likely represented dMCs **(Figure 2N,O,Q)**. By 7 dpp, nearly all *atoh1a+* cells were newly derived (**Figure 2P,Q)**. Based on these results, we concluded that skin injury triggers *de novo* production of dMCs and MCs within the trunk epidermis and that early stages of epidermal regeneration are associated with a high proportion of dMCs. Thus, dMCs are a transient *atoh1a+* cell type during both skin development and regeneration.

### dMCs exhibit molecular features of both basal keratinocytes and MCs

To characterize the molecular and cellular properties of regenerating dMCs, we focused on 5 dpp, a timepoint that contains a mix of dMCs and MCs (**Figure 2K**). We first assayed expression of Tp63, a transcription factor critical in epidermal development and highly expressed in basal keratinocytes (Bakkers et al., 2002; Guzman et al., 2013; Lee and Kimelman, 2002; Rangel-Huerta et al., 2021) (**Figure 3B, asterisks**). Immunostaining revealed that MCs did not express detectable levels of Tp63 (**Figure 3A-A’**). By contrast, dMCs exhibited significantly higher levels of Tp63 staining than MCs, albeit at lower levels than basal keratinocytes, with ∼75% of dMCs staining positive for Tp63 (**Figure 3B-C**). Thus, dMCs express Tp63 at intermediate levels relative to basal keratinocytes and MCs.

**Figure 3.**
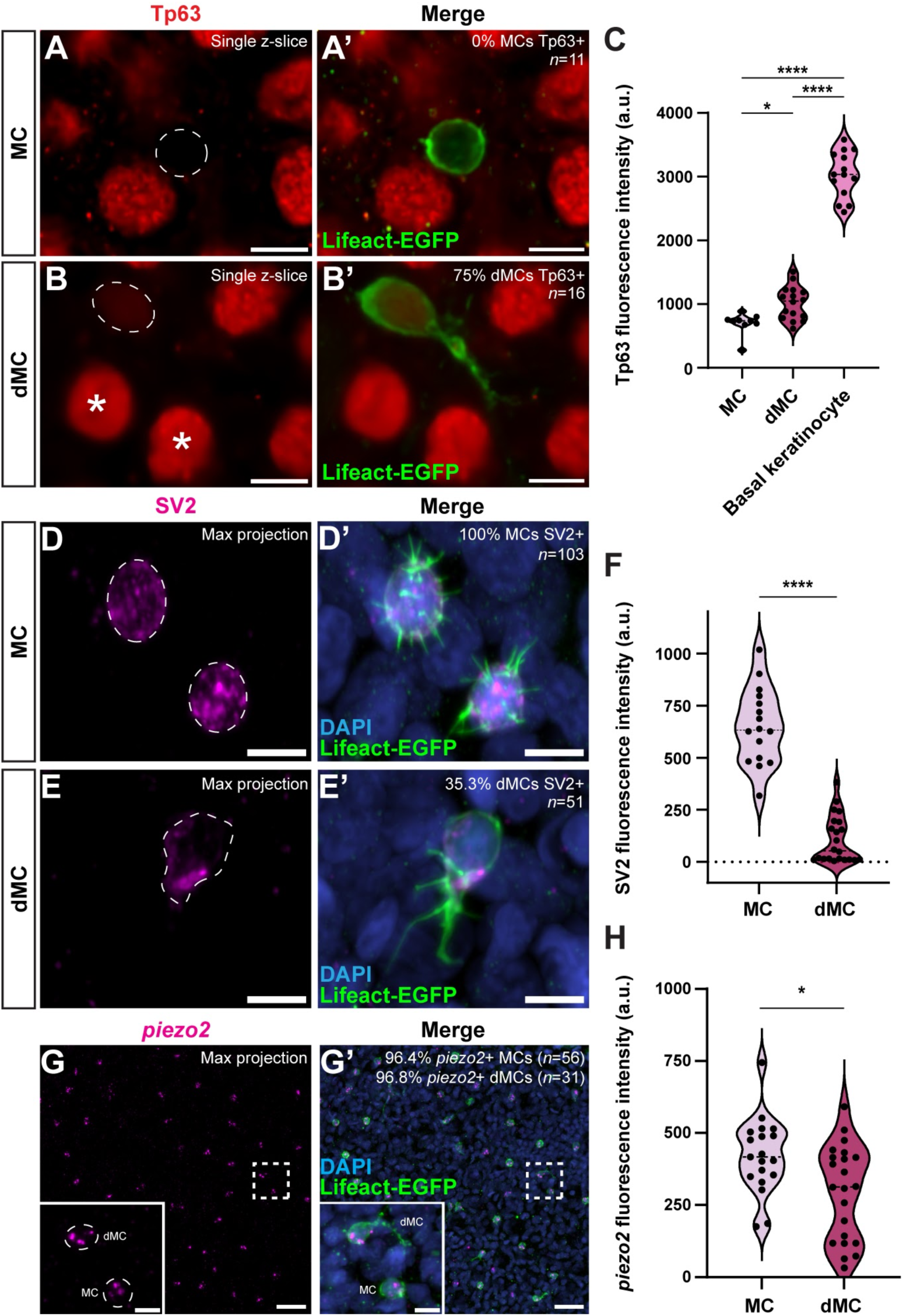
dMCs share molecular features with keratinocytes and MCs. (**A-B)** Representative micrographs of MCs and dMCs within the 5 dpp regenerating scale epidermis of a *Tg(atoh1a:lifeact-egfp)* adult visualized with anti-Tp63 (red) and anti-GFP (green) antibodies. Mature MCs are Tp63 negative (A) whereas dMCs are often weakly Tp63 positive (B). Dashed lines outline the nuclei. Asterisks indicate basal keratinocyte nuclei. **(C)** Violin plots of Tp63 staining intensity in MCs, dMCs, and basal keratinocytes. Confocal micrographs acquired with identical imaging parameters were analyzed for Corrected Total Cell Fluorescence (CTCF) measured in arbitrary units. Each dot represents an individual cell (*n*=9 MCs, *n*=16 dMCs, *n*=14 basal keratinocytes). A one-way ANOVA with post-hoc Tukey HSD test was used to compare Tp63 staining levels between cell types. **(D-E)** Representative micrographs showing MCs and dMCs within the 5 dpp regenerating scale epidermis of a *Tg(atoh1a:lifeact-egfp)* adult stained with anti-SV2 (magenta) and anti-GFP (green) antibodies. MCs are always SV2 positive (D), whereas some dMCs (E) are SV2 positive. Dashed lines outline the cell bodies. **(F)** Violin plots of SV2 staining intensity quantified using CTCF in MCs and dMCs. Each dot represents an individual cell (*n*=16 MCs, *n*=22 dMCs). A non-parametric Mann-Whitney test was used to compare SV2 staining levels between cell types. **(G)** Representative micrographs of MCs and dMCS within the 5 dpp regenerating scale epidermis of a *Tg(atoh1a:lifeact-egfp)* adult stained with an anti-*piezo2* HCR probeset (magenta) and an anti-GFP (green) antibody. Dashed box indicates region magnified in inset. Both MCs and dMCs are *piezo2* positive. **(H)** Violin plots of *piezo2* HCR staining intensity in MCs and dMCs quantified using CTCF. Each dot represents an individual cell (*n*=19 MCs, *n*=22 dMCs). A non-parametric Mann-Whitney test was used to compare *piezo2* staining levels between cell types. *, P<0.05; ****, P<0.0001. Scale bars, 5 µm (A-B,D-E,G inset) and 20 µm (G).

Given our finding that dMCs expressed *atoh1a*, we postulated that dMCs might share additional molecular properties with MCs. MCs express markers of neuroendocrine and mechanosensory function, such as synaptic vesicle glycoprotein 2 (SV2) and the mechanosensitive cation channel Piezo2, respectively. Consistent with previous observations during ontogeny (Brown et al., 2023), we found that regenerated MCs expressed SV2 (**Figure 3D-D’**). SV2 staining of dMCs in regenerating skin revealed a mix of SV2+ and SV2– dMCs (**Figure 3E-E’**). We quantified the SV2 fluorescence intensity from dMCs with detectable signal and found dMCs displayed lower SV2 levels than MCs (**Figure 3F**). Hybridization chain reaction (HCR) staining for *piezo2* labeled both MCs (**Figure 3G-G’, arrowhead**) and dMCs (**Figure 3G-G’, arrow**) in regenerating skin. Analysis of the *piezo2* signal revealed slightly lower levels in dMCs compared to MCs (**Figure 3H**). In summary, regenerating dMCs display molecular properties that overlap with both basal keratinocytes and MCs, suggesting they may represent a transitional or immature MC.

### MC and dMC protrusions are dynamic and differentially polarized

Apart from previous reports that tracked MC turnover over the course of days or weeks by imaging cytoplasmic or nuclear reporters (Brown et al., 2023; Wright et al., 2017), little is known about the *in vivo* dynamics of MCs. Given the differences in cell shapes between dMCs and MCs during development and regeneration, we sought to compare the dynamics and polarities of the actin cytoskeleton in these two populations. To this end, we mounted and intubated *Tg(atoh1a:lifeact-egfp)* animals and performed live-cell confocal microscopy of fully intact skin over several hours. Imaging of MCs in adult zebrafish revealed strong Lifeact-EGFP signal in microvilli as expected (**Figure 4A, arrowhead**). We captured numerous microvillar reorientation, retraction, and extension events occurring in all MCs (**Figure 4A-A’’’’; Movie 1**), suggesting previously unappreciated MC microvillar dynamics. MC microvilli rarely extended >3 microns from the membrane and often appeared to coalesce or merge together, although we lacked the resolution to reliably characterize these merging events. Despite these microvillar dynamics, the MC cortex remained spherical during imaging (**Figure 4A-A’’’’**). By analyzing reconstructed cross-sections of MC microvilli in animals also expressing a Cdh1 (E-cadherin) knock-in reporter to label keratinocyte membranes [Cdh1-tdTomato; (Cronan et al., 2018)] (**Figure 4C**), we found the majority of MC microvilli extended in basal keratinocyte-facing or lateral directions, whereas periderm-facing processes were less common (**Figure 4D,D’,F,G**).

**Figure 4.**
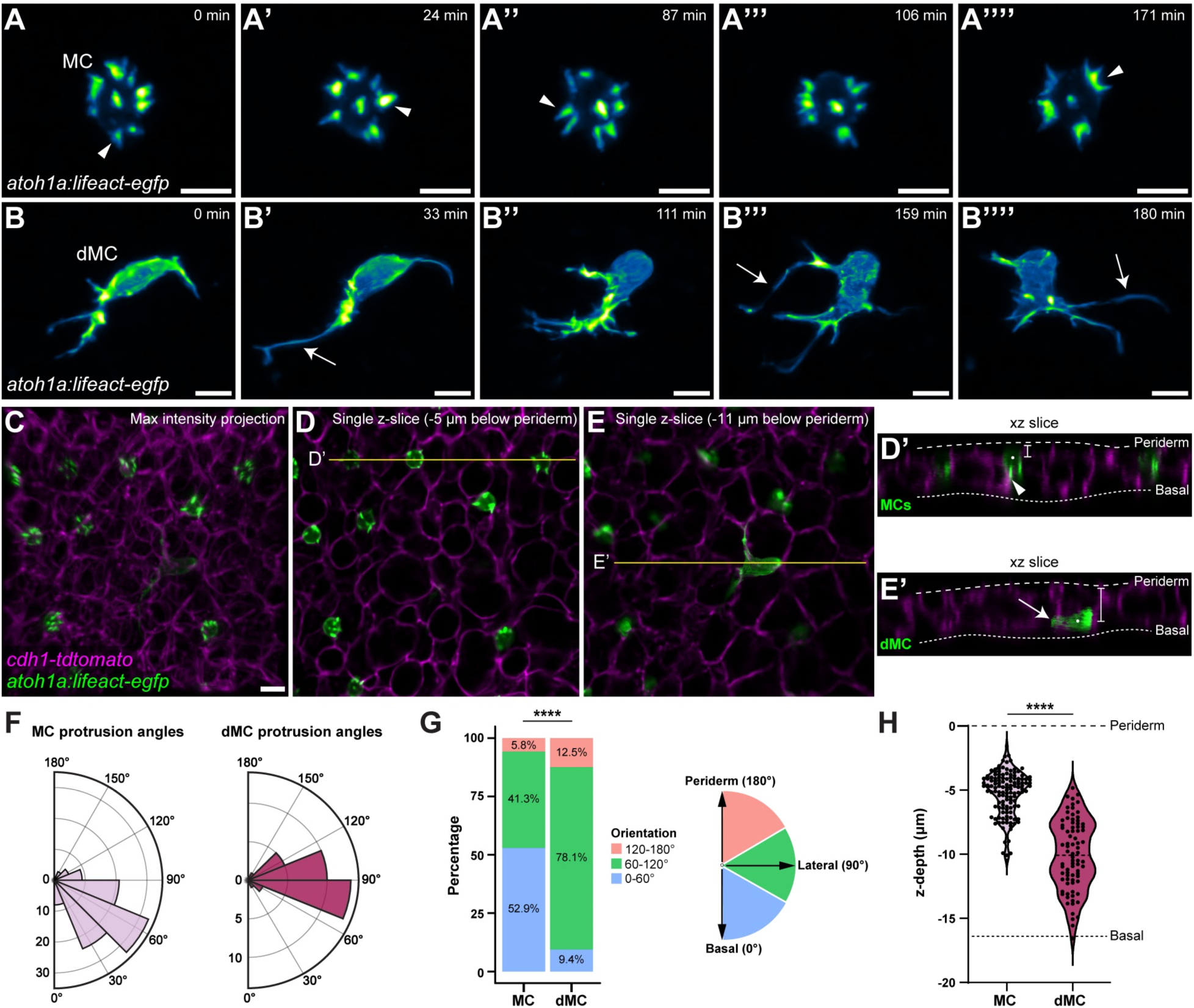
MC and dMC protrusions are dynamic and differentially polarized in the skin. **(A-A’’’’)** Still images of a timelapse of a single MC expressing *Tg(atoh1a:lifeact-egfp)*. White arrowheads point to microvilli extension, retraction, or merging events. **(B-B’’’’)** Still images of a timelapse of a single dMC visualized with *Tg(atoh1a:lifeact-egfp)*, image acquisition as described in (A). Note the longer, filopodial-like protrusions (white arrows) and highly amorphous cell body. **(C-E)** Representative maximum intensity projection (C) or individual z-slices (D,E) of the scale epidermis in an animal expressing *Tg(atoh1a:lifeact-egfp)* to label dMCs and MCs (green) and *(cdh1-tdTomato)* to label lateral cell membranes (magenta). **(D’,E’)** Reconstructed xz slices along the yellow lines in D and E. Dashed lines indicate the outer and inner epidermal margins. Note that the MCs in D’ are located near the upper, periderm layer of the skin with basal facing protrusions (arrowhead), whereas the dMC in E’ is located near the lower, basal layer of the skin with a lateral facing protrusion (arrow). Brightness and contrast were adjusted in (E,E’) to better illustrate the dMC morphology. **(F)** Polar histograms of MC and dMC protrusion angles. Lifeact-EGFP+ protrusions that could be individually resolved in 3D were measured relative to the z-axis of the epidermis using the coordinate system diagrammed in (G). Shaded regions indicate the number of protrusions falling in that angle range. Plots show *n*=104 protrusions from 10 MCs and *n*=32 protrusions from 11 dMCs. **(G)** Stacked bar charts depicting results in (F), with protrusions binned based on orientation. A χ^2^ test was used to compare between cell types (****, P<0.0001; χ^2^ statistic, 18.9903). **(H)** Violin plots of z-depths of MCs and dMCs measured relative to the periderm surface in the scale epidermis of *n*=130 MCs and *n*=80 dMCs from 5 fish (11-12 mm SL). The average depth of the basal surface of basal keratinocytes in the data set is indicated by a horizontal dashed line (−16.4 µm). A non-parametric Mann-Whitney test was used to compare between cell types (****, P<0.0001). Scale bars, 5 µm (A-C).

In contrast to MC microvilli, imaging of dMCs in juveniles revealed long, thin Lifeact-EGFP+ protrusions that routinely exceeded 10 microns in length and in extreme cases exceeded 25 microns (**Figure 4B-B’’’’, arrowheads**). dMCs dynamically rearranged both their protrusions and cell cortex (**Figure 4B-B’’’’; Movie 2**). dMC protrusions tended to coalesce and extend from one end of the cell, with the opposite side of the dMC having few or no protrusions (**Figure 4B’’**). The majority of dMC protrusions extended in lateral or periderm-facing directions, whereas basal keratinocyte-facing processes were less common (**Figure 4E,E’,F,G).** Reconstructed cross-sections also revealed that MCs tended to reside just below the periderm, whereas dMCs were commonly in lower strata (**Figure 4D’,E’,H**). Thus, our observations indicate MCs and dMCs occupy different strata of the epidermis and utilize dynamic actin-based protrusions that are of distinct sizes and polarities.

### dMCs are mesenchymal-like cells that directly mature into MCs

We wondered how MCs and dMCs behaved during periods of development when MC density greatly increases in the trunk skin (10-15 mm SL, **Figure 1J,K**). Based on their association with somatosensory neurites and desmosome-like contacts with keratinocytes, we hypothesized that MCs would be relatively immotile. By contrast, our Cre lineage tracing data (**Figure 1O**) and observations of dMC protrusion dynamics and polarity (**Figure 4**), led us to hypothesize that dMCs may migrate within the epidermis.

To characterize MC and dMC motility, we intubated *Tg(atoh1a:lifeact-egfp)* juveniles and imaged fields of view containing MCs and dMCs over 6 h (**Figure 5A, Movie 3**). Consistent with our hypotheses, cell tracking revealed that MCs were largely immotile, whereas dMCs migrated laterally within the epidermis (**Figure 5A, insets)**. dMC motility was highly variable: On average, dMCs migrated 2.4 µm/hr during imaging, but some moved upwards of 8 µm/hr, while others remained stationary (**Figure 5B**). By contrast, MCs were immotile, with an average displacement of 0.8 µm/hr, which likely reflects imaging drift (**Figure 5B**). dMCs were most motile when adopting elongated, ovoid cell bodies with long, unipolar protrusions at one end of the cell (**Figure 5E, pink arrowheads**). By contrast, dMCs with multipolar protrusions were largely immotile (**Figure 5E, orange arrowheads**). Although the majority of dMCs traveled in a persistent or linear manner (**Figure 5C**), dMCs had a significantly reduced track displacement compared to total distance traveled (**Figure 5D**), suggesting that dMCs migrate directionally but often switch directions. Consistent with this notion, dMC-dMC or dMC-MC contacts resulted in lateral dMC movement away from the contact (**Movie 4**). Together these observations suggest that MCs are immotile, epithelial-like cells, whereas dMCs are motile, mesenchymal-like cells that undergo contact inhibition upon encountering another *atoh1a+* cell.

**Figure 5.**
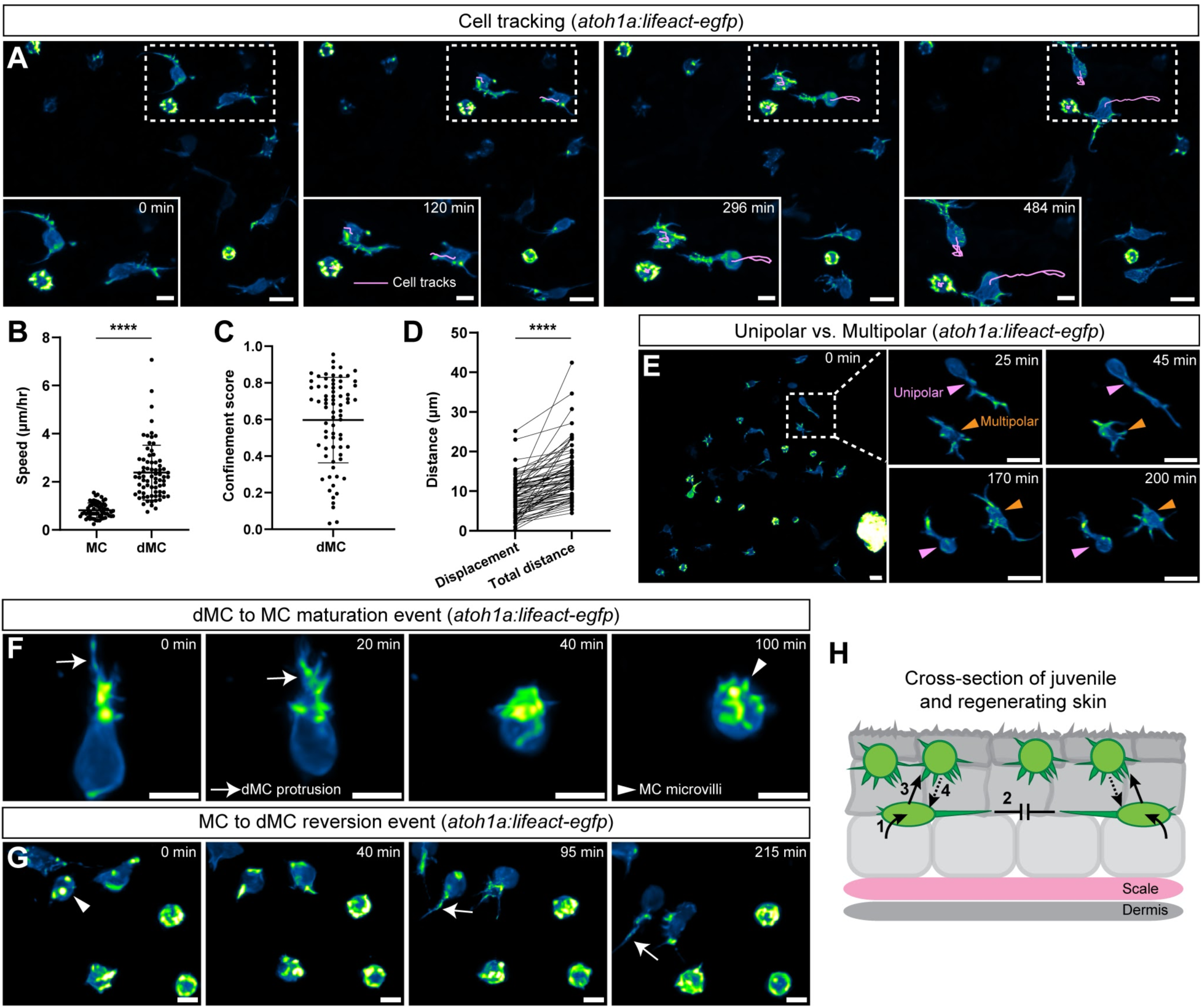
dMCs are motile cells that can directly mature into MCs in developing zebrafish skin. **(A)** Cell tracks (magenta) of individual MCs and dMCs traced over time. **(B)** Dot plot of cell speed (µm/hour) of individually tracked cells (*n*=60 MCs and *n*=74 dMCs). A non-parametric Mann-Whitney test was used to compare speed between cell types (****, P<0.0001). **(C)** Cell tracks scored for confinement ratio where values near 0 are indicative of confined movement and values near 1 are indicative of a linear trajectory (*n*=74 dMCs). **(D)** Dot plot of dMC track displacement, which measures the distance between the starting and ending point of each cell track, and total distance traveled (*n*=74 dMCs). A paired Mann-Whitney test was used to compare displacement and distance (****, P<0.0001). In (B-D), horizontal lines indicate the mean and error bars indicate the SD. **(E)** Representative timelapse stills from *Tg(atoh1a:lifeact-egfp)* expressing juvenile zebrafish skin. Magenta arrowheads highlight dMCs in their migratory configuration with unipolar protrusions. Orange arrowheads highlight dMCs in their exploratory or immotile configuration with multipolar protrusions. Note that individual cells can switch between the unipolar and multipolar configurations. **(F)** Representative timelapse stills of a dMC to MC maturation event. Arrow indicates dMC protrusion retraction and arrowhead indicates formation of microvilli. Note the cell body switches from an ovoid to spherical shape during this transition. **(G)** Representative timelapse stills of a MC to dMC reversion event. Arrowhead indicates microvilli and arrow indicates filopodial-like protrusion extension. **(H)** Schematic depicting proposed model of dMC maturation events described in this study. (1) dMCs emerge in lower epidermal strata from *ΔNp63+* basal keratinocyte progenitors; (2) dMCs are motile cells that migrate in the direction of their protrusions and can undergo contact inhibition upon encountering another dMC; (3) dMCs can directly adopt the mature MC morphology in upper epidermal strata; and (4) in rare instances, MCs can revert to a dMC morphology. Scale bars, 10 µm (A), 5 µm (A, insets), 10 µm (E, E, insets), and 5 µm (F,G).

MCs are generally considered post-mitotic (Merot and Saurat, 1988; Moll et al., 1996; Vaigot et al., 1987; Weber et al., 2023; Wright et al., 2017). Indeed, we did not observe instances of MC cell division in our live-imaging of either homeostatic juvenile or regenerating adult skin. By contrast, dMCs divided in both contexts (**Figure S1A-C**). During cell division, dMCs resorbed their protrusions, developed a smooth actin-rich membrane, and underwent cytokinesis to generate two daughter dMCs, which established protrusions orthogonal to the division plane and migrated away from each other (**Figure S1A-C; Movie 5)**. Due to its experimental tractability, we quantified dMC and MC cell division frequency during skin regeneration. To this end, we injected 5-ethynyl-2’-deoxyuridine (EdU), a thymidine analog incorporated into DNA during active DNA synthesis, at 4 dpp and fixed and stained scales at 5 dpp (**Figure S1D**). In this experimental paradigm, no MCs stained positive for EdU, whereas 65% of dMCs stained positive for EdU (**Figure S1E,F**). Thus, in addition to exhibiting differences in motility, dMCs and MCs show contrasting cell cycle states.

Our findings that dMCs exhibited characteristics of both basal keratinocytes and MCs (**Figure 3**) suggested they may represent an intermediate cellular state between these two cell types. Indeed, during our live-imaging of juvenile and regenerating adult skin, we observed a small subset of dMCs (*n*>10) withdraw their long protrusions, round up their cell body, and rapidly extend microvilli reminiscent of the mature “mace-like” MC morphology (**Figure 5F; Movies 6,7**). These observations are consistent with the direct maturation of a dMC into a MC. In rare instances (*n*=2), we observed the reciprocal process, i.e., the reversion of a MC into a dMC-like morphology (**Figure 5G; Movie 8**), suggesting that some MCs can switch between cell states. Thus, based on these observations, we conclude that dMCs are capable of cell migration, cell division, and direct maturation into MCs (**Figure 5H**).

### Genetic loss of Eda results in altered dMC and MC morphologies in trunk skin

Eda signaling promotes the development of diverse vertebrate skin appendages (reviewed by Sadier et al., 2014), including zebrafish scales (Harris et al., 2008). We previously reported that zebrafish homozygous for a presumptive null *eda* allele (*eda^dt1261/dt1261^,* hereafter *eda^-/-^*) displayed reduced MC density within the trunk, but not facial, epidermis (Brown et al., 2023).

We postulated that Eda signaling may also promote the morphological maturation of MCs. To address this hypothesis, we incrossed *eda^+/-^* adults and used *Tg(atoh1a:lifeact-egfp)* localization to compare MC morphology between homozygous mutants and sibling controls. As expected, control trunk MCs developed a typical “mace-like” morphology decorated by microvilli (**Figure 6A-A’’’,G**). By contrast, we observed a striking and highly penetrant loss of MC microvillar structures along the trunk of *eda^-/-^* mutants (**Figure 6B-B’’’,G**). In *eda^-/-^* trunk MCs, Lifeact-EGFP signal accumulated near the membrane in a ring-like fashion (**Figure 6B’, arrowhead**), reminiscent of the morphology of dMCs during cell division (**Figure S1A-C**). The few MC microvilli that formed in *eda^-/-^* trunk MCs were short and thin, making them difficult to resolve with confocal microscopy. To determine whether Eda globally regulated MC morphology throughout the skin, we imaged MCs in the corneal epidermis, which is not squamated. This analysis revealed corneal MCs with similar microvilli-rich morphologies in *eda^-/-^*mutants and siblings (**Figure 6C-D,G**). Finally, to determine whether loss of Eda signaling also impacted dMC morphology, we compared the maximum protrusion length of dMCs in *eda^-/-^* mutants and controls and found that mutant dMCs had significantly longer protrusions (**Figure 6E,F,H;** *eda^-/-^* dMC mean lengths = 16.3 μm, siblings = 8.7 μm). In extreme cases, dMC protrusions reached almost 40 microns in length in *eda^-/-^* trunk epidermis (**Figure 6F**). Together, these results indicate that Eda is necessary for the normal morphologies of dMCs and MCs specifically within the trunk skin compartment.

**Figure 6.**
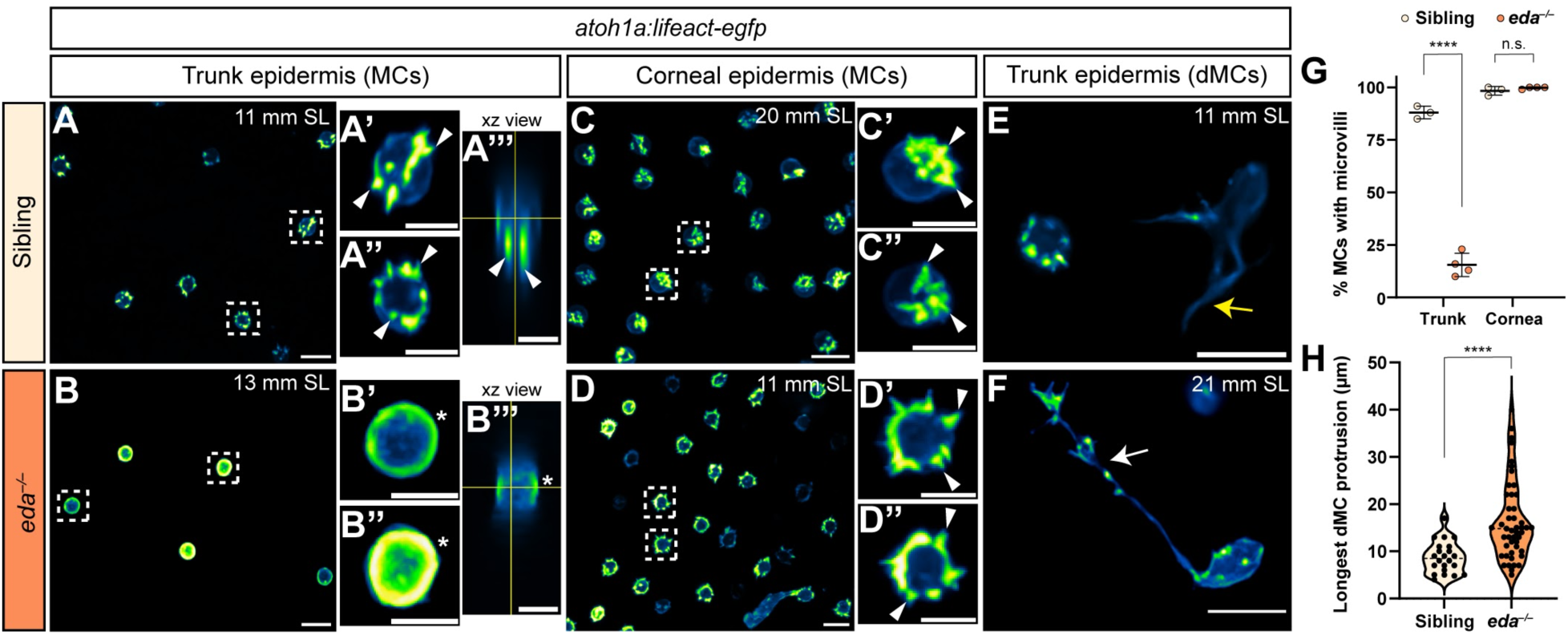
Genetic loss of Eda results in altered dMC and MC morphologies in trunk skin. **(A,B)** Representative confocal images of MCs within juvenile trunk epidermis of animals of the indicated genotypes and stages. Dashed boxes indicate cells magnified in (A’,A’’,B’,B’’). Arrowheads indicate microvilli and asterisks indicate intense Lifeact-EGFP signal forming a smooth, ring-like cortical structure evident in cross section in the *eda^-/-^* mutant epidermis. **(C,D)** Representative confocal images of MCs within corneal epidermis of animals of the indicated genotypes. Dashed boxes indicate cells magnified in (C’,C’’,D’,D’’). Arrowheads indicate microvilli. **(E,F)** Representative confocal images of dMCs within the trunk epidermis of animals of the indicated genotypes and stages. Yellow arrow in E indicates the longest protrusion on a dMC in *eda* sibling epidermis. White arrow in F indicates the longest protrusion on a dMC in *eda^-/-^* mutant epidermis. **(G)** Quantification of percentage of MCs with discernable microvilli in trunk or corneal epidermis of animals of the indicated genotypes. Each dot indicates a juvenile (9-20 mm SL) zebrafish of the respective genotype where a collection of images were analyzed (sibling trunk, *n*=386 cells from 3 fish; *eda^-/-^* trunk, *n*=235 cells from 4 fish; sibling cornea: *n*=169 cells from 4 fish; *eda^-/-^* cornea, *n*=332 cells from 4 fish). Fisher’s exact test shows a significant difference between genotypes in the trunk but not cornea (****, P<0.0001; n.s., P=0.3413). **(H)** Violin plots of the longest dMC Lifeact-EGFP+ protrusion within the trunk epidermis of juveniles of the indicated genotypes. Each dot represents an individual cell (siblings, *n*=22 dMCs; *eda^-/-^*, *n*=47 dMCs). A non-parametric Mann-Whitney test (****, P<0.0001) was used to compare between cell types. Scales bars, 10 µm (A-F), and 5 µm (A’-A’’’, B’-B’’’, C’-C’’, D’-D’’).

## DISCUSSION

The defining morphological features of MCs allows for their identification across diverse types of vertebrate skin (Hartschuh et al., 1986; Whitear, 1989). How MCs acquire this precise morphology during skin development or regeneration is unknown. While basal keratinocytes have been identified as MC progenitors in mammalian and zebrafish skin, intermediate steps in the MC lineage progression remain poorly understood. Previous histology-based experiments noted transitional cells that could serve as cellular intermediates during MC maturation (English, 1974; English, 1977; Garant et al., 1980; Tachibana and Ishizeki, 1981; Tachibana and Nawa, 1980). However, these studies lacked the temporal resolution to rigorously establish a relationship between transitional cells and MCs. Here, we directly address this knowledge gap by using the optical accessibility of zebrafish skin to track the emergence of MCs during development and regeneration. Our results support a model in which dMCs emerge in lower epidermal strata and migrate locally within the epidermis. Their migration occurs both laterally, presumably due to their lateral-facing actin-rich membrane protrusions, and vertically within the epidermis. As they mature in superficial strata, dMCs reorganize their actin cytoskeleton to elaborate polarized microvilli, thereby adopting the terminal mace-like morphology characteristic of MCs (**Figure 5H**).

### A transient cell state during development and regeneration

We observed that dMCs appeared transiently during a period of skin organogenesis associated with dermal appendage growth along the trunk. dMC frequency peaked during a time of rapid increase in MC density. These findings closely mirror previous studies in other vertebrates. For instance, the “dendritic:globular” MC ratio sharply decreased in embryonic and human fetal palmar skin prior to the peak of MC density (Kim and Holbrook, 1995). Furthermore, our lineage tracing identified basal keratinocytes as dMC precursors, supporting previous ultrastructural findings that suggested basal keratinocytes give rise to transitional cells (English, 1974; English, 1977; Garant et al., 1980; Tachibana and Ishizeki, 1981; Tachibana and Nawa, 1980). However, since those TEM studies did not reconstruct transitional cell morphology, any relationship between transitional cells and dMCs has remained unclear. Based on our lineage tracing and morphological data, we propose that dMCs and transitional cells likely represent similar cellular intermediates. However, dMCs may represent just one of several intermediate states between basal keratinocytes and MCs, each with their own characteristics as previously suggested (Tachibana and Ishizeki, 1981).

Severe skin wounds, such as burns, can alter sensory perception, thereby degrading patient quality of life (reviewed by Girard et al., 2017). Human MC regeneration after injury appears poor (Stella et al., 1994), although engineered skin grafts present a promising avenue (Hahn et al., 2019). Studies of the mechanisms of MC regeneration in model organisms may inform improved wound healing treatments. Previous work in mammalian models found that MCs regenerate after peripheral nerve crush (Burgess et al., 1974; Nurse et al., 1984), cauterization of touch domes (Horch, 1982), full thickness wounds (Tachibana and Ishizeki, 1981), or repeated skin shaving (Weber et al., 2023; Wright et al., 2017). We found that scale plucking induced a high proportion of dMCs during the early phase of scale regeneration and that MC density ultimately recovered to pre-injury levels. Using a photoconvertible reporter, we established that MCs in regenerated epidermis form predominantly due to *de novo* regeneration rather than migration from surrounding epidermal reservoirs. Due to the rapid kinetics, scale pluck presents a tractable model for reliably triggering dMC/MC production after cutaneous injury.

We used scale pluck as an approach to reliably induce dMCs and quantitatively characterize their molecular properties during regeneration. We found that dMCs expressed the basal keratinocyte marker Tp63 at intermediate levels relative to MCs (low) and basal cells (high). We found that dMCs frequently expressed the MC markers *piezo2* and SV2, albeit at levels lower than MCs. Together these data lend further support to the notion that transitional cells and dMCs are one and the same, and suggest that dMCs may represent MC precursors. Additionally, *piezo2* expression in dMCs suggests they may have the capacity for mechanosensory function. Examining whether mechanosensation impacts any of the dMC behaviors described here is an exciting area for future study.

### Dynamic cellular behaviors of dMCs and MCs

The microvilli of several cell types, including cultured renal and immune cells, exhibit dynamic motility (Cai et al., 2017; Meenderink et al., 2019). However, to our knowledge, no live-cell imaging of MC microvilli has been reported. Our *in vivo* imaging revealed that MC microvilli were dynamic and frequently extended, rearranged, or retracted. Our observations are consistent with the lack of desmosomes along MC microvilli, a feature noted in a wide range of vertebrates and skin compartments (Düring and Andres, 1976; Garant et al., 1980; Halata, 1975; Hashimoto, 1972; Iggo and Muir, 1969; Landmann and Halata, 1980; Takahashi-Iwanaga and Abe, 2001). We further found that MC microvilli were polarized in lateral- or basal keratinocyte-facing orientations. Intriguingly, lamprey microvilli face both superficial and basal strata (Daghfous et al., 2020; Takahashi-Iwanaga and Abe, 2001), whereas MC microvilli within cat touch domes face superficial strata (Iggo and Muir, 1969). These observations suggest malleability of MC microvilli polarity across vertebrates. If microvilli are involved in MC function as previously proposed (Garant et al., 1980; Takahashi-Iwanaga, 2003; Takahashi-Iwanaga and Abe, 2001; Toyoshima et al., 1998; Yamashita et al., 1993), reconciling microvillar dynamics and polarity with MC function will be an interesting area for future exploration.

Our live imaging further revealed a number of previously unrecognized behaviors of dMCs, many of which are properties of mesenchymal cells. First, we found that dMCs elaborate long, thin actin-rich protrusions, reminiscent of protrusions previously observed with cytokeratin 20 staining in gerbil palate (Tachibana et al., 1997). These protrusions extended from lateral-facing membranes and interdigitated between basal and suprabasal keratinocytes. Second, in contrast to the immotile MCs, we observed that dMCs migrated laterally within the epidermis. dMC migration correlated with the presence of unipolar protrusions, whereas dMCs with multipolar protrusions were largely stationary. Intriguingly, dMCs with similar shapes (termed unipolar and globular) were previously noted in adult human oral mucosa (Tachibana et al., 1998). Future work is needed to dissect the molecular composition of dMC protrusions to determine whether they are similar to filopodial/filopodial-like protrusions observed in other contexts. Third, we observed that dMC-dMC or dMC-MC contacts resulted in repulsive behaviors, akin to contact inhibition. As zebrafish MCs are exclusively solitary cells, we speculate that this behavior may promote cell dispersion. Fourth, we observed that dMCs could divide, generating two daughter cells with similar morphologies and behaviors. This suggests that at least a subset of dMCs may be transient amplifying cells. Finally, and perhaps most significantly, we observed in rare instances dMCs could mature into MCs by retracting their protrusions, adopting a spherical cell body, and elaborating microvilli. This provides the first direct evidence that dMCs can serve as MC precursors. Although we do not know yet what cues this maturation, it appears to involve a transition from a mesenchymal-like to an epithelial-like state.

### Ectodysplasin and MC maturation

In contrast to the well-studied stereocilia of hair cells (reviewed by Barr-Gillespie, 2015), virtually nothing is known about molecular control of MC microvilli assembly. Nevertheless, both hair cell stereocilia and MC microvilli contain a core of parallel actin bundles and share expression of at least one actin bundling protein (Sekerková et al., 2004; Sekerková et al., 2006). We previously found that Eda signaling was partially necessary for MC development along the trunk skin and that *eda* mutants had a reduced rate of MC production (Brown et al., 2023). Here, we extended these studies by examining the morphology of dMCs and MCs in *eda* loss of function mutants. Surprisingly, we found that most MCs completely lacked microvilli in *eda* mutants, a novel MC phenotype to our knowledge. This observation indicates Eda functions upstream of a MC microvillar program; dissecting the connections between Eda signaling, MC microvilli, and MC function represent important areas for further study, and *eda* mutants offer a promising system to test MC sensory ability in the absence of microvilli. Furthermore, we found that dMCs developed hyperextended protrusions in *eda* mutants. One interpretation of these data is that dMC protrusion depends on local dMC and/or MC density. Addressing this possibility will require rigorous spatial analysis and experimental manipulation independent of Eda signaling.

In summary, we use the live imaging advantages of zebrafish to describe the dynamic cell behaviors of dMCs and MCs during development and regeneration. We establish that dMCs represent an intermediate cell state between basal keratinocytes and MCs. We further identify Eda signaling as essential for the regional morphogenesis of MCs. Finally, we speculate that the mesenchymal-like behaviors of dMCs may have relevance for understanding the genesis of Merkel cell carcinoma.

## MATERIALS AND METHODS

### Animals

#### Zebrafish and developmental staging

Zebrafish were housed at 26-27℃ on a 14/10 h light cycle. Zebrafish of either sex were used. Strains used: AB (Wild-Type), *(cdh1-tdTomato)^xt18^* (Cronan et al., 2018)*, Tg(atoh1a:nls-Eos)^w214Tg^* (Pickett et al., 2018)*, Tg(atoh1a:lifeact-EGFP)^w259Tg^, TgBAC(ΔNp63:Cre-ERT2)^w267Tg^* (Brown et al., 2023)*, Tg(actb2:LOXP-BFP-LOXP-DsRed)^sd27Tg^* (Kobayashi et al., 2014)*, Tg(Ola.Sp7:mCherry-Eco.NfsB)^pd46Tg^* [referred to as *Tg(sp7:mCherry)*] (Singh et al., 2012), and *eda^dt1261^* (Harris et al., 2008). All zebrafish experiments were approved by the Institutional Animal Care and Use Committee at the University of Washington (protocol #4439-01).

To control for differences in growth rates, zebrafish post-embryonic development was staged based on standard length (SL) (Parichy et al., 2009). SL of fish was measured using the IC Measure software (The Imaging Source) on images captured on a Stemi 508 stereoscope (Zeiss) equipped with a DFK 33UX264 camera (The Imaging Source). *eda* mutants and siblings were sorted by visible phenotype starting at 7 mm SL. Mutants were grown separately from siblings.

#### Induction of TgBAC(ΔNp63:Cre-ERT2) with 4-OHT

To activate recombination with Cre-ERT2, 1 dpf *TgBAC(ΔNp63:Cre-ERT2)*; *Tg(actb2:LOXP-BFP-LOXP-DsRed); Tg(atoh1a:lifeact-EGFP)* embryos were treated with 10 μM 4-OHT for 24 hr and screened for successful recombination at 3-5 dpf as evidenced by DsRed+ epidermal cells. 4-OHT was prepared as described (Felker et al., 2016).

*Scale pluck*.

A scale pluck protocol was used either to collect scales for immunofluorescence or to induce scale regeneration. To prepare fish for scale pluck, fish were anesthetized in 0.006-0.012% buffered MS-222 (MilliporeSigma, Cat #: E10521) diluted in system water until a surgical plane of anesthesia was reached. Anesthetized fish were moved under a dissecting microscope and placed on the lid of a petri dish. Dumont #5 forceps were used to pluck scales from the trunk in a posterior to anterior procession. Fish were then returned to system water and monitored for full recovery.

### Imaging

#### Scale regeneration time course

Scales were plucked from adult *Tg(atoh1a:lifeact-egfp);Tg(sp7:mcherry)* zebrafish as described above. To facilitate subsequent imaging, the two rows along the midline were plucked. Plucked fish were returned to the recirculating system and imaged using confocal microscopy at the indicated dpp. 1.5x and 8x images were collected for quantification and downstream analyses.

#### Whole animal photoconversion

Prior to scale removal, *Tg(atoh1a:nls-Eos)* zebrafish were exposed to light from a UV LED flashlight (McDoer) for 15 min in a reflective chamber constructed from a styrofoam box lined with aluminum foil. A similar lateral region of the trunk was imaged over subsequent days identified by approximate body position below the dorsal fin and relative to underlying pigment stripes.

#### Confocal imaging

Short-term live imaging of juvenile and adult zebrafish was achieved by anesthetizing fish in 0.006-0.012% MS-222 diluted in system water for ∼5 min. Once the fish was immobilized, it was mounted in a custom imaging chamber and the fish body was secured by carefully adding molten 1% agarose in system water. Agarose was not applied to the trunk region where imaging occurred or near the gills and mouth of the fish to ensure survival. Agarose embedded fish were then covered with MS-222 solution and placed under an A1R MP+ scanhead mounted on a Ni-E upright microscope (Nikon). Except where noted otherwise, a 16x water dipping objective (N.A. 0.8) was used for z-stack acquisition and images were post-processed using the Denoise.ai function in NIS-Elements (Nikon). For short-term imaging, fish were taken off the microscope after a maximum 25 min of image collection and returned to system water for recovery.

Long-term live imaging of juvenile zebrafish was achieved through an intubation-based protocol that delivered tricaine-water to immobilized zebrafish using a peristaltic-pump, similar to Xu et al. (2015). In brief, fish were anesthetized in MS-222 for 8-10 min until gill movement became very slow. Fish were transferred to a custom imaging chamber and embedded with 1% agarose as described above. Continuous delivery of MS-222 was achieved by using forceps to gently insert polyethylene tubing (Becton Dickinson #427421 or #427400) into the fish’s mouth. This delivery line was held in place by modeling clay and 0.08% MS-222 was delivered to the fish by a peristaltic pump at a flow rate of 2-3 ml per min. Multipoint time lapse imaging was achieved through Nikon elements software by manually setting large z-stacks for each field of view to account for sample drift during imaging. Depending on the experiment, time lapse images were acquired every 3-5 mins for up to 6 h at room temperature. To revive fish after multi-hour time lapse imaging, the peristaltic pump line was transferred to a bottle containing system water and fish were closely monitored for recovery. Data acquired from zebrafish that did not survive the intubation session were excluded from further analysis.

### Staining

#### Alizarin Red S staining

Alizarin Red S stains calcium deposits allowing visualization of osteoblast-derived zebrafish scales. To visualize zebrafish scale development, live animals were stained for 20 min in a solution of 0.01% (wt/vol) Alizarin Red S (ACROS Organics, Cat #: 400480250) dissolved in system water and shielded from light, rinsed 3x, 5 mins in system water, then transferred back into fresh system water as described (Bensimon-Brito et al., 2016).

#### Immunofluorescence

Zebrafish were anesthetized in a solution of 0.012% MS-222 in system water for 2 min. Approximately 30 scales were plucked and transferred to 1.5 mL tubes containing 375 μl 1xPBS. For fixation, 125 μl of 16% paraformaldehyde (PFA) was added to achieve a final 4% PFA solution. Scales were incubated for 20 min at room temperature on a gently rotating platform. PFA was washed out by carefully removing the fixation solution and replacing it with 400 μl of 0.2% PBST (1xPBS + 0.2% Triton X-100). Scales were washed 3x, 5 min with PBST then blocked with blocking solution (10% normal goat serum in PBST) for 2-3 h at room temperature. Blocking solution was removed, and 200 μl of primary antibody solution made up in blocking solution was added. Primary antibodies used: Mouse monoclonal anti-SV2 (DSHB Cat #: SV2, RRID:AB_2315387) at 1:50; or Rabbit polyclonal anti-GFP (Thermo Fisher Scientific, Cat #: A11122, RRID:AB_221569) at 1:500. Scales were incubated in primary antibody overnight at 4℃, protected from light. The next day, scales were washed 4x, 15 min with PBST before adding secondary antibody made up in blocking solution. Secondary antibodies used: Goat anti-Mouse Alexa Fluor 647 (Thermo Fisher Scientific, Cat #: A32728, RRID:AB_2633277) at 1:500; or Goat anti-Rabbit Alexa Fluor 488 (Thermo Fisher Scientific, Cat #: A32731, RRID:AB_2633280) at 1:1000. Scales were incubated for 2 h at room temperature, protected from light. Secondary solution was washed out 4x, 15 min using PBST. To visualize nuclei, 5 ng/μl DAPI (MilliporeSigma, Cat #: 508741) was added. DAPI was washed out 4x, 5 min using PBST. Scales were mounted epidermis-side up between a microscope slide and coverslip in ProLong Gold (Thermo Fisher Scientific, Cat #: P36930). Imaging was performed with a 40x (NA 1.3) oil immersion objective.

#### Hybridization chain reaction

HCR on adult zebrafish scales using a custom *piezo2* probe set (Molecular Instruments; accession #: XM_021468270.1; set size: 20; amplifier: B3) was performed as described (Brown et al., 2023). Scales were imaged with a 25x water immersion objective (N.A. 1.1)

#### EdU labeling during scale regeneration

Scales were plucked from adult *Tg(atoh1a:lifeact-egfp);Tg(sp7:mCherry)* zebrafish to induce scale regeneration. Fish were returned to their tanks for recovery prior to EdU injection. Regenerating fish were anesthetized in MS-222 and placed ventral side up on a sponge situated under a dissecting microscope. 10 μl of 10 mM EdU (Thermo Fisher Scientific, Cat #: C10640) was injected intraperitoneally in the region between the pelvic fins. Fish were transferred back to system water for recovery. Approximately 20 h later, these fish had their regenerating scales plucked and subjected to EdU staining following the manufacturer’s instructions. If downstream immunofluorescence was required, this was performed after EdU detection. Stained scales were imaged with a 40x (NA 1.3) oil immersion objective.

### Image and statistical analysis

#### Image processing

Image processing was performed using FIJI/ImageJ (Schindelin et al., 2012) or Imaris (Oxford Instruments). Unless otherwise indicated, all images were gathered through z-stack acquisition and displayed as maximum intensity projections. Throughout the figures, single channel images of *Tg(atoh1a:lifeact-egfp)* are color-coded according to signal intensity 564 using the “Green Fire Blue” lookup table in FIJI/ImageJ.

#### Cell counting & shape analysis

*atoh1a+* cells were classified as dMCs or MCs as follows: Confocal z-stacks were max projected and the ImageJ “Cell Counter” function was used to manually classify *atoh1a+* cells as dMC or MC based on cell morphology. For shape analysis, max projected images were thresholded using Huang’s algorithm. Using the “Analyze Particles…” function, circularity and roundness were calculated. In Figure 1N, “circular” cells were defined as those having circularity and roundness values >0.7.

#### Fluorescence intensity analysis

Fluorescence intensity of MCs and dMCs was calculated from confocal z-stacks acquired using identical settings by Corrected Total Cell Fluorescence (CTCF) in ImageJ. CTCF = Integrated Density – (Area of selected cell X Mean fluorescence of background readings).

#### z-depth analysis

For the z-depth analysis in Figure 5H, confocal z-stacks were acquired using a 25x water dipping objective (N.A. 1.1) and a z-step of 0.3 μm. The outer surface of the periderm was defined as a depth of 0. The distance from the center of the cell bodies of dMCs and MCs to the periderm surface was measured on reconstructed yz slices using the line tool in ImageJ.

#### Cell tracking

To quantify cell motility metrics, the trackmate plugin in ImageJ was used. z-stack acquired timelapse images were imported and made into max projections. To correct drift encountered during long term imaging acquisition, the Imagej plugin “correct 3D drift” was used to correct xy drift. Trackmate was opened and manual detection was performed by tracking individual cells of interest. The metrics calculated from trackmate (Total distance traveled, confinement ratio, etc) are explained https://imagej.net/plugins/trackmate/algorithms.

#### Statistical analysis

Statistical analysis and graphing were performed using R (R Core Team, 2023), MATLAB (MathWorks) or GraphPad (Prism). Statistical tests used and sample numbers are described in the corresponding figure legend.

## ACKNOWLEDGEMENTS

We thank the LSB Aquatics staff for animal care and Dr. David Tobin for sharing zebrafish. The authors are grateful to all members of the Rasmussen lab for discussion, technical assistance, and support. The SV2 monoclonal antibody used in this study was obtained from the Developmental Studies Hybridoma Bank, created by the NICHD of the NIH and maintained at The University of Iowa, Department of Biology, Iowa City, IA 52242.

## AUTHOR CONTRIBUTIONS

**Evan W. Craig**: Conceptualization, Investigation, Methodology, Visualization, Writing - Original Draft, Funding acquisition; **Erik C. Black**: Investigation, Methodology; **Camille E.A. Goo**: Investigation, Methodology; **Avery Angell Swearer**: Investigation, Methodology; **Nathaniel G. Yee**: Investigation, Methodology; **Jeffrey P. Rasmussen**: Conceptualization, Investigation, Methodology, Validation, Supervision, Visualization, Writing - Original Draft, Funding acquisition.

## DATA AVAILABILITY

All data needed to evaluate the conclusions in the paper are present in the paper and/or the Supplementary Materials.

## SUPPLEMENTAL MOVIE LEGENDS

**Movie 1. Microvilli dynamics in zebrafish MCs.** Timelapse movie of MCs expressing *Tg(atoh1a:lifeact-egfp)* in juvenile zebrafish skin. Note spike-like microvilli that extend, retract, and coalesce. Scale bar, 5 µm.

**Movie 2. Actin dynamics in zebrafish dMCs.** Timelapse movie of a dMC expressing *Tg(atoh1a:lifeact-egfp)* in juvenile zebrafish skin. Note thread-like filopodial protrusions that are highly dynamic. Scale bar, 10 µm.

**Movie 3. Long-term imaging reveals dMC motile behaviors.** 6 hour timelapse movie of MCs and dMCs expressing *Tg(atoh1a:lifeact-egfp)* in an intubated juvenile zebrafish. Manual cell tracks (white traces) show dMCs crawling in the direction of their protrusions while MCs remain largely stationary. Scale bar, 10 µm.

**Movie 4. dMC-dMC contact inhibition.** Timelapse of a dMC-dMC contact (indicated by arrow at time 0 min) that results in both cells migrating in opposing directions. Scale bars, 10 µm.

**Movie 5. *In vivo* dMC cell division in juvenile zebrafish skin.** Timelapse movie of a dMC expressing *Tg(atoh1a:lifeact-egfp)* undergoing cell division. In all dMC division events recorded, dividing dMCs retract a single unipolar protrusion, form a circular and actin-rich membrane, and undergo cytokinesis. Daughter cells then rapidly extend filopodial protrusions and migrate opposite from one another. Scale bar, 10 µm.

**Movie 6. dMC to MC maturation event.** Timelapse of single dMC expressing *Tg(atoh1a:lifeact-egfp)* that withdraws its filopodial protrusions, rounds its cell cortex, and extends small microvillar extensions as observed in mature MCs. Scale bar, 5 µm.

**Movie 7. dMC behaviors during regeneration**. Timelapse of the regenerating scale epidermis at 3 dpp in a *Tg(atoh1a:lifeact-egfp)* zebrafish. Labels indicate different dMC behaviors including maturation, motility, and cell death. Scale bar, 10 µm.

**Movie 8. MC to dMC reversion event.** Timelapse of developing *Tg(atoh1a:lifeact-egfp)* expressing zebrafish skin with a mix of MCs and dMCs in the field of view. Note a cell with a MC morphology begins to transition into a dMC-like cell with a long unipolar protrusion. Scale bar, 10 µm.

## SUPPLEMENTAL FIGURE AND LEGEND

**Figure S1:**
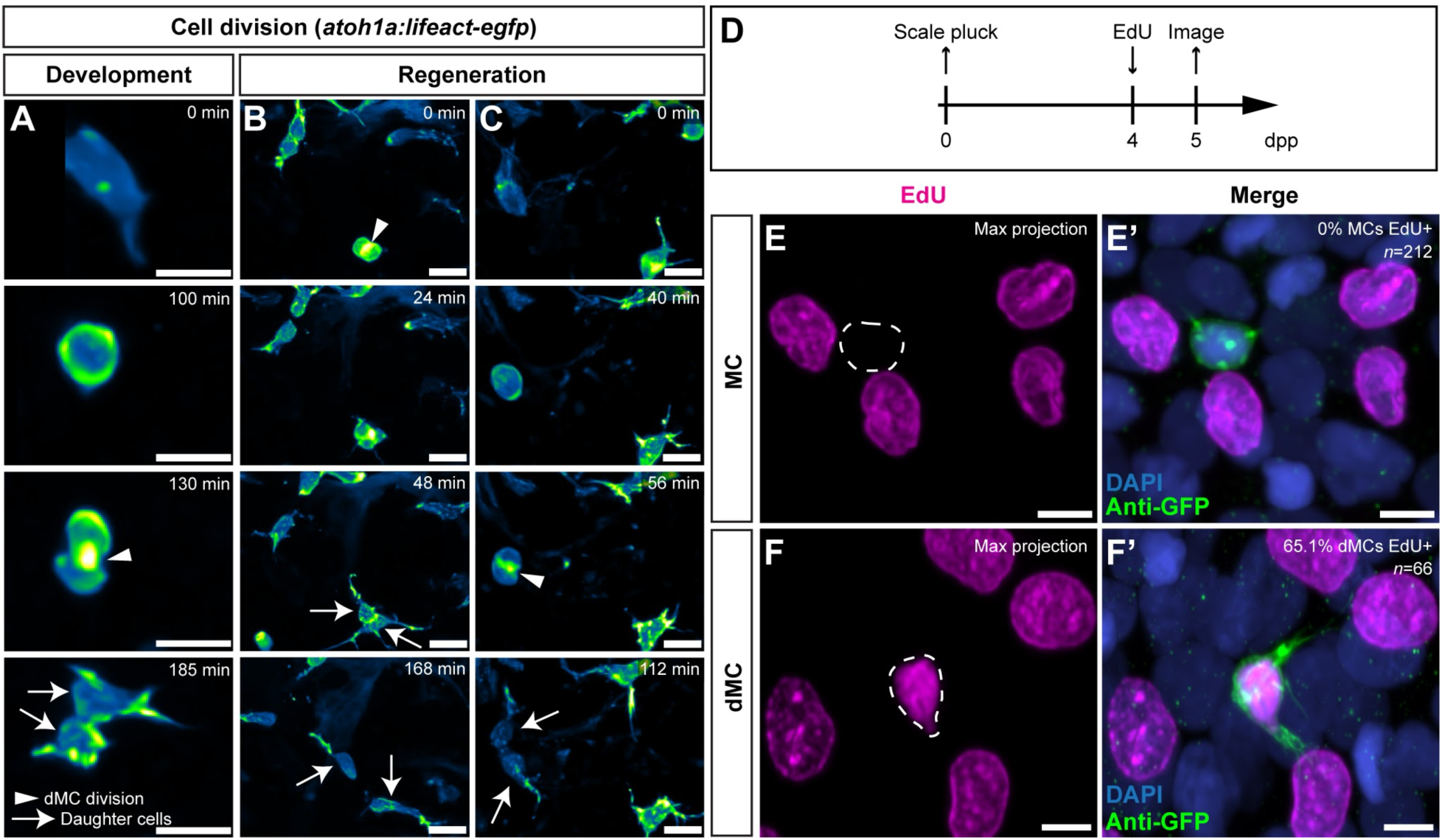
Differences in cell cycle state of dMCs and MCs. **(A-C)** Still images from timelapses showing cell divisions of dMCs expressing *Tg(atoh1a:lifeact-egfp)* in the juvenile (A) or regenerating scale epidermis (B,C). Arrowheads denote cytokinetic furrows and arrows denote daughter cells. **(D)** Experimental design of EdU administration and imaging during scale regeneration. **(E,F)** Representative confocal images of EdU and anti-GFP staining in regenerating *Tg(atoh1a:lifeact-egfp)* scales. White dashed lines indicate nuclear outlines. Scale bars, 10 µm (A-C) and 5 µm (E-F).

